# Maltodextrin administration ameliorates brain pathology in a mouse model of mitochondrial disease

**DOI:** 10.1101/2023.06.28.546916

**Authors:** Adán Domínguez-Martínez, Esther Molina-Menor, Marcos Blanco-Ramos, Andrea Urpi, Juli Peretó, Manuel Porcar, Albert Quintana

## Abstract

Mitochondrial dysfunction lead to a wide group of progressive and fatal pathologies known as mitochondrial diseases (MD). One of the most common pediatric representation of MD is Leigh Syndrome, affecting 1/40.000 births. LS is characterized by neurodegeneration in specific brain areas, such as brainstem and basal ganglia, and by respiratory and motor alterations. However, the results obtained from clinical trials based on antioxidant therapies are controversial. Thus, the development novel antioxidant strategy is required to improve the efficacy of current palliative treatments. In this regard, Ndufs4KO mouse model is a suitable model to test new drugs in the field of MD and LS. Therefore, we set to assess the therapeutic potential of oral administration of *Micrococcus luteus*, a high-antioxidant content microorganism. Incidentally, we identified that while *M. luteus* administration did not possess any beneficial actions, the cryopreservant maltodextrin (MDX), included in the preparation, ameliorated the phenotype of Ndufs4KO mice. Our results show that MDX treatment at a concentration of 30 g/L increased lifespan and reduced microglial reaction compared to vehicle-treated Ndufs4KO mice. However, no improvement in locomotion nor respiratory function was observed in MDX-treated mice compared to vehicle-treated Ndufs4KO mice. Metataxonomic characterization of intestinal microbiome identified differential profiles in Ndufs4KO mice at the genus level. Furthermore, MDX treatment increased the variability of the abundance of *Akkermansia sp*. Thus, this work paves the way for further studies to confirm the therapeutic potential of MDX in mitochondrial disease.

## Introduction

Mitochondria are organelles that play a central role in homeostasis and survival as they provide the energy required for proper cell function (Friedman & Nunnari, 2014). Mutations that impair normal mitochondrial function have a huge impact on organs with high energy requirement, such as the brain or muscles, and lead to a wide group of progressive and often fatal pathologies known as primary mitochondrial diseases (MD) (Ng & Turnbull, 2016; Vafai & Mootha, 2012).

MD have a prevalence of 1/5000 newborns, being Leigh Syndrome (LS) the most common pediatric presentation (1/40000 births). LS onset is typically described between 3 and 12 months of age, although onset in prenatal and adult life has also been reported, and is commonly caused by mitochondrial complex I deficiency (almost 50 % patients) (Gerards et al., 2016). However, over 75 mitochondrial DNA (mtDNA) and nuclear DNA (nDNA) mutations have been described to cause LS (Lake et al., 2016). Despite the genetic and clinical heterogeneity, LS patients commonly present symmetrical brain lesions, mainly in the brainstem and basal ganglia, together with failure to thrive, persisting vomiting, hypotonia, motor deficits and respiratory failure (Arii & Tanabe, 2000; Rahman, 2012; Sofou et al., 2014).

Unfortunately, the severity of the disease, the variability in its progression and the paucity of good animal models difficult and limit our understanding of the pathophysiology of LS. Moreover, current treatment for LS and other MD are limited and palliative, thus urging the need to develop novel drugs to cope with the symptomatology (Lake et al., 2016). In this regard, the characterization of a mouse model lacking the *Ndufs4* gene (Ndufs4KO), essential for the mitochondrial complex I of the respiratory chain, represented an important step forward in the study of the disease (Kruse et al., 2008; Quintana et al., 2010).

Ndufs4KO mice faithfully recapitulate human LS condition, being, thus, suitable for testing new drugs and therapeutic approaches (Bolea et al., 2019). These animals show increased oxidative stress (OS) and reactive oxygen species (ROS) levels in affected brain areas, leading to neurodegeneration and glial activation (Angelova & Abramov, 2018; Liu et al., 2015; Quintana et al., 2010). OS causes lipid peroxidation and neuron membrane alteration, DNA damage (especially mitochondrial DNA damage), protein oxidation and subsequent alteration of signaling pathways (Cobb & Cole, 2015; Grimm & Eckert, 2017; Lavie, 2015). Moreover, these high OS and ROS levels seem to be responsible for the symptomatology of MD and LS given the observed susceptibility of the nervous system to OS damage (Ng & Turnbull, 2016; Vafai & Mootha, 2012). Furthermore, the model also shows motor alterations, respiratory deficits, epilepsy and premature death and has revealed the prominent role of the dorsal brainstem in disease progression and lethality (Bolea et al., 2019; Quintana et al., 2012). Given the role of OS and ROS in MD, antioxidants have proven a good palliative treatment to ameliorate the clinical progression of LS in animal models in preclinical assays (Liu et al., 2015). Even though the role of antioxidant treatments in clinical trials remains unclear due to failure of several approaches in improving clinical progression (Parkinson et al., 2013), it has been described that administration of the antioxidant EPI-743 ameliorated symptomatology of LS pediatric patients (Martinelli et al., 2012). For this reason, novel antioxidant-based strategies are still of interest in the field of LS treatment (Enns, 2014).

In the last decades, the role of microbiota in health and disease has been in the spotlight of many scientists. The microbiota can be defined as the microbial population inhabiting a specific environment at a specific time, and particularly gut microbiota has received interest since differences between healthy and ill patients of a group of diseases have been found. It has been suggested that microbes inhabiting the intestinal tract of animals, including humans, may contribute to the global health state of the individual and, therefore, to the appearance and progression of certain pathologies (e.g., neurologic, respiratory, autoimmune and metabolic disorders). However, whether changes in microbiota are cause or result of these altered states remains a question to be answered in most cases (Fan & Pedersen, 2021; Lynch & Pedersen, 2016).

Specifically, microorganisms inhabiting the intestinal tract can modulate OS levels and the antioxidant capacity of the host in the central nervous system. Thus, their contribution to the development of several neurodegenerative disorders is being increasingly recognized (Dumitrescu et al., 2018). In the era of pre, pro and postbiotics (live biotherapeutic microorganisms), an approach that deeply studies the effect of potentially new probiotic microorganisms or specific substances on the microbiota, and further in the general health state of the individuals is of interest (Cunningham et al., 2021; Soheili et al., 2022).

In this regard, carotenoids are pigments present in many phototrophic and non-phototrophic organisms (including vegetables, algae and bacteria) with widely-known antioxidant properties (Milani et al., 2017). In fact, microbial communities from extremophilic and highly irradiated environments (e.g., solar panels, the supralittoral area of the coast or deserts) are enriched in carotenoid-synthetizing microorganisms with biotechnological potential in biomedicine given their antioxidant capacity (Asker et al., 2007; Molina-Menor et al., 2020; Molina-Menor et al., 2019; Tanner et al., 2019; Tapia et al., 2021; Tian & Hua, 2010). These antioxidant properties are of interest since ROS molecules are generated not only during aerobic metabolism, but also during pathological processes in degenerative diseases and under abiotic stress conditions (Apel & Hirt, 2004; Stahl & Sies, 2005).

We hypothesized that antioxidant microorganisms and/or microbial extracts may have a therapeutic effect on MD. Therefore, we set to develop a complete strategy to characterize the effect of carotenoid-rich microbial formulations, containing freeze-dried microorganisms and maltodextrin (MDX), on the phenotype of Ndufs4KO. Our results indicate that while no significant effect of microorganisms per se was observed, the administration of maltodextrin leads to an amelioration of clinical and histological signs in Ndufs4KO, which may be mediated by an effect on microbiota profiles.

## Materials and Methods

### Animal model

*Ndufs4*^*Δ/Δ*^ (Ndufs4KO) mouse were generated by Dr.Palmiter lab (Kruse et al., 2008) and characterized by Quintana et al., (2010) were used in this study. All mice were on a C57BL6/6J background after backcrossing for at least 10 generations. Deletion of the exon 2 in *Ndufs4* gene was confirmed by PCR. Female and male mice of the same age were used and maintained on a rodent diet (5053, Picolab) and water available *ad libitum*, in an animal facility at 22 °C with a 12 h light-dark cycle. All procedures were approved by the animal care committees at the Universitat Autònoma de Barcelona and Generalitat de Catalunya.

### Physiological and behavioral parameters

Body weight and gait/postural alterations were measured every day for each mouse. Mice were euthanized after losing 20 % of their maximum weight.

### Maltodextrin dose-response study

Low-DE maltodextrin (MDX) (Mane Iberica S.A) was diluted in drinking water at different concentrations (1.5, 5, 15 and 30 g/L), and provided to the animals *ad libitum*. Water was used as vehicle (VEH, no treatment). MDX beverage was replaced twice a week to avoid drink contamination by fungi. MDX treatment started after weaning at P21. Given the severe locomotion alterations of Ndufs4KO mice, MDX beverage was poured over chao pellets (one per animal) and placed at the same location in the cage daily.

### Behavioral assays

Behavioral assays were performed at early-stage (P30-35), mid-stage (P40-45) and late-state (P50-55). Phenotypic markers (weight, hydration and survival) were also registered and analyzed as stated below:

|  | Behavioral tests |  |  | Phenotypic markers |  |  |
| --- | --- | --- | --- | --- | --- | --- |
|  | Rotarod | Plethysmography | Open Field | Weight | Hydration | Ataxia<br>Curling<br>Twisting |
| Age<br>(days) | P30<br>P40<br>P50 | P32<br>P42<br>P52 | P35<br>P45<br>P55 | Daily | Daily | Daily |

#### Rotarod test

A rotarod device was used to assess motor coordination and global physical condition of animals as described (Bolea et al., 2019). Mice were placed on a gradually accelerating rotating drum (from 4 rpm to 50 rpm) for a maximum of 3 min. Each trial ended when the mouse fell off the rod. The assay consisted in 3 trials, separated by 5 min rest period.

#### Open-field

Mice were placed in the center of an open-field arena (650 × 365 × 400 mm) and allowed to move freely for 45 min in order to assess locomotor activity as described (Bolea et al., 2019). Travelled distance and velocity average were analyzed in two periods of time: the first 5 min, and the rest 40 min. The time spent inside and outside the center of the cage was also analyzed. EthoVision tracking software (Noldus) was used for analysis.

#### Whole-body plethysmography

Ventilatory function was assessed through whole-body plethysmography under unrestrained conditions as described (Prada-Dacasa et al., 2020). Mice were placed in an airtight chamber that allowed free movement and spontaneous breathing. The system was calibrated to 1 mL. Mice were monitored for 1 h: 45 min of adaptation and 15 min of measurement. Respiratory frequency (RF; breaths/min) and tidal volume normalized per body weight (TV; μL/g) were measured.

### Tissue preparation and immunofluorescence

Brains were dissected after euthanasia, fixed overnight in 4 % paraformaldehyde (PFA) in PBS (pH 7.4) and then cryopreserved in a PBS solution with 30 % sucrose (w/v) as described (Quintana et al., 2010). Mouse brains were sectioned at 30 μm in a cryostat and stored in PBS solution for further use. Brain sections containing the VN were obtained from bregma -5.6 to bregma -6.96. For immunofluorescence, brain sections were rinsed in 0.2 % Triton X-100 in PBS (PBST). An incubation step of 60 min at room temperature (RT) was done with a solution of 10 % Normal Donkey Serum (NDS) in PBST to block non-specific binding. The sections were then incubated overnight with primary antibodies αGFAP Ch (ref PA1-10004) 1:2000 and αIba1 Rb (ref 019-19741) 1:2000 at 4 °C. Brain sections were further washed in PBST and incubated for 1 h at RT with secondary antibodies αCh 488 (green) 1:500 and αRb 594 (red) 1:500 in 1% NDS-PBST, and finally washed with PBS and rinsed in water before mounting onto glass slides for microscopic analysis.

### High-throughput 16S rRNA gene sequencing (metataxonomy)

DNA extractions were carried out with the Stool DNA Isolation Kit Dx from NORGEN Biotek (Ontario, Canada) following the manufacturer’s instructions. Gut samples in PBS 1X were first centrifuged for 12 min at 12000 rpm at room temperature. The pellets were then resuspended in 400 μL of PBS 1X, which were further used for the DNA extraction. The conserved regions V3 and V4 of the bacterial 16S rRNA gene were amplified with primers 5′-TCG TCG GCA GCG TCA GAT GTG TAT AAG AGA CAG CCT ACG GGN GGC WGC AG 3′ and 5′-GTC TCG TGG GCT CGG AGA TGT GTA TAA GAG ACA GGA CTA CHV GGG TAT CTA ATC C -3’, respectively. Libraries were constructed following the Illumina’s standard protocol and Illumina MiSeq platform (2×300 pb) was used for amplicon sequencing. Bioinformatic analysis was carried out by Darwin Bioprospecting Excellence S.L.

### Statistics

GraphPad v8.0 software was used for statistical analysis. The tests were chosen according to the experimental design, as stated in figure legends. Statistical significance is also stated in the legend. Data are shown as the mean±SEM.

## Results

### Maltodextrin administration extends survival in Ndufs4KO mice

To assess the potential role of a microorganism-based antioxidant administration we delivered an extract of *Micrococcus luteus*, a microorganism we had previously identified to possess high antioxidant properties (Molina-Menor et al., 2019), to a cohort of Ndufs4KO mice. Similarly to LS patients (Sofou et al., 2014), Ndufs4KO mice present a severe phenotype including death at early stage (around week 7) (Quintana et al., 2012). Administration of *M. luteus* mixed in the drinking water (5 g/L) led to a significant extension of the lifespan of Nduf4KO compared to our historical record for Ndufs4KO mice (median survival of 69 vs 43 days, respectively (**Figure 1A**). However, similar effects were obtained when a cohort of Ndufs4KO mice receiving 5g/L maltodextrin, used as a cryopreservant in the preparation of the *M. luteus* compared to a vehicle (water)-treated Ndufs4KO mice (**Figure 1B**).

**Figure 1.**
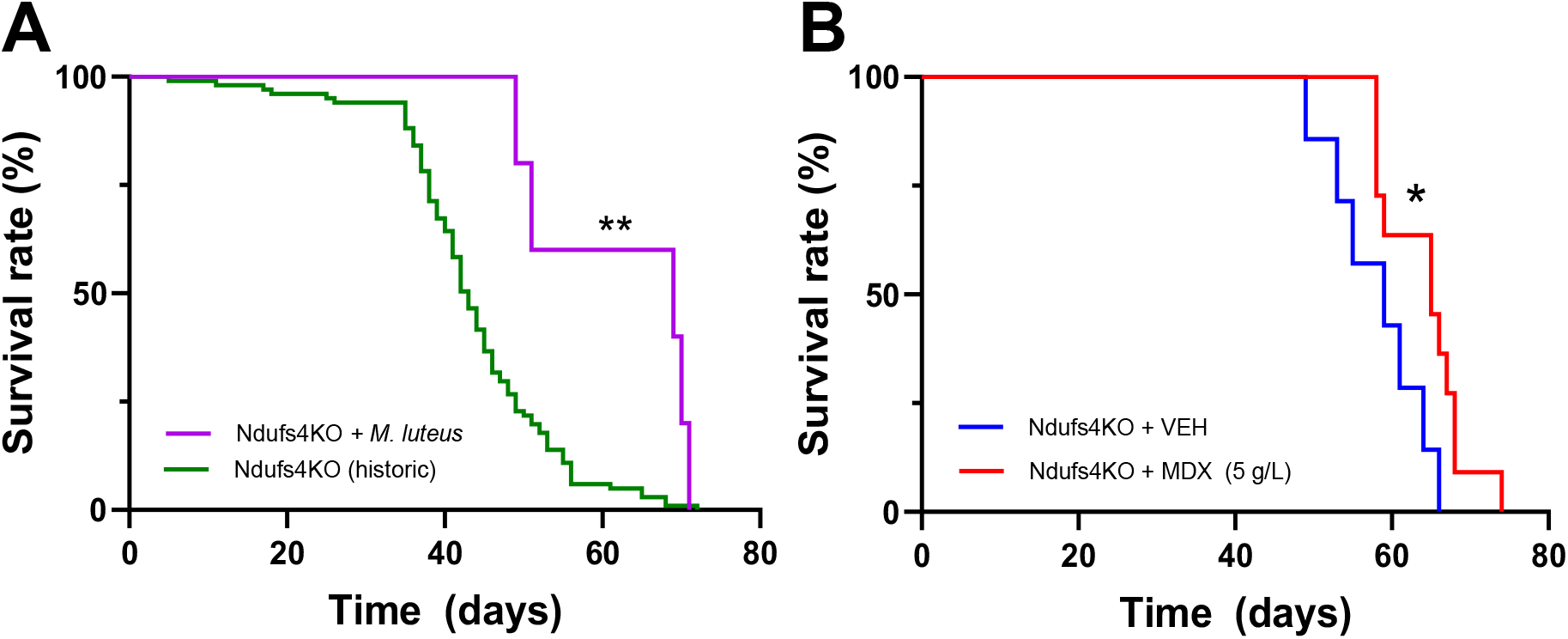
Maltodextrin administration extends lifespan in Ndufs4KO mice. **A)** Survival curve of Ndufs4KO mice receiving a 5 g/L solution of *Micrococcus luteus* and maltodextrin (n=5) in the drinking water vs. a comparative historical survival of the Ndufs4KO colony (n=101). **p<0.01 vs. Ndufs4KO historical record, Long-rank test. **B)** Survival curve of Ndufs4KO mice receiving a 5 g/L solution of maltodextrin in the drinking water (MDX, n=11) vs. mice receiving regular drinking water (VEH, n=7). *p<0.05 vs. VEH, Long-rank test.

To determine if the lifespan survival of Ndufs4KO mice is dependent on MDX dosage, a dose-response study was designed. To that end, we administered increasing concentrations of MDX (1.5, 5, 15 and 30 g/L) or vehicle (water). Subsequently a log-rank test was performed to analyze the effect of different doses of MDX on the survival of Ndufs4KO mice. Our results showed that MDX significantly increased survival in a compared to the vehicle-treated group, obtaining a significant lifespan extension in Ndufs4KO mice treated with 30 g/L and 5 g/L of MDX (**Figure 2**). Median survivals in the MDX 5 g/L and 30 g/L groups were 66 and 67 days, respectively, significantly higher than in vehicle-treated group, which showed a median rate of survival of 55 days. Noteworthy, no effect was observed at an intermediate dose of 15 g/L, even though an increase in median survival was found (56 vs. 60, veh vs 15 g/L, respectively).

**Figure 2.**
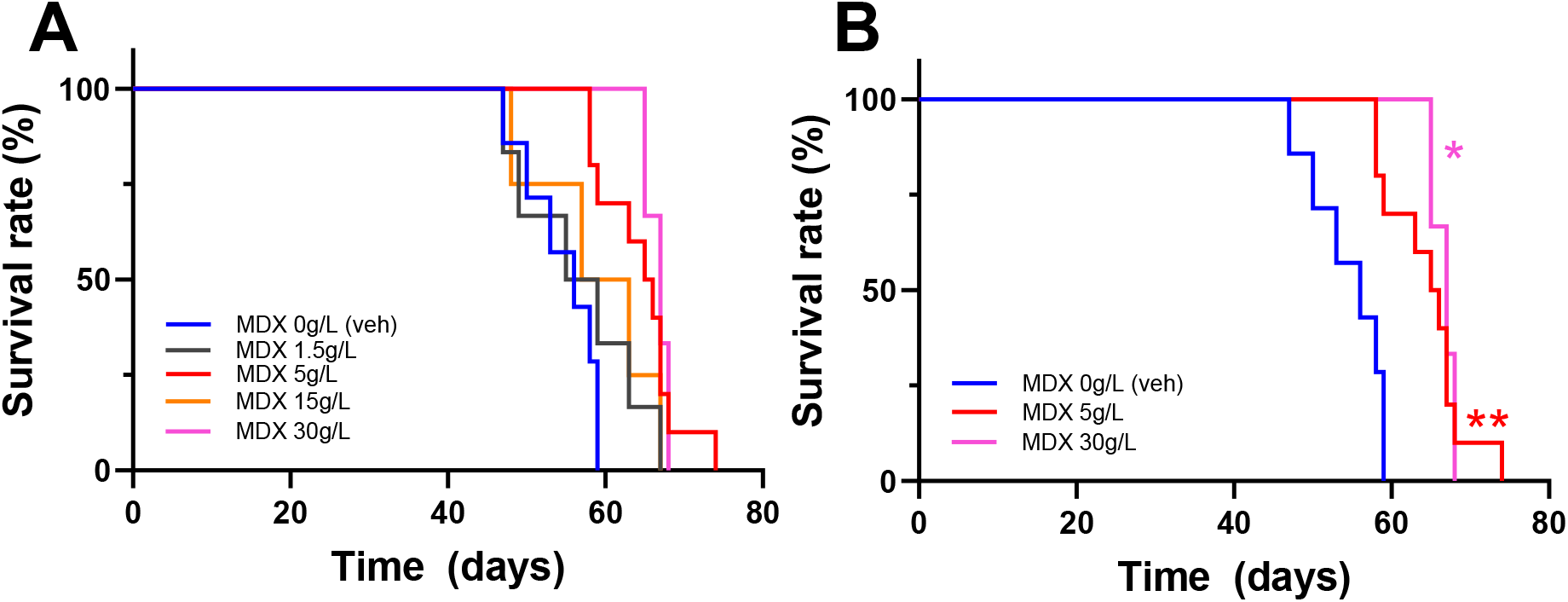
Effect of increasing maltodextrin concentrations on Ndufs4KO survival. **A)** Combined survival curves of 0-30 g/L MDX doses. (0 g/L -veh-n=7, 1.5 g/L n=6, 5 g/L n=10, 15 g/L n=4, 30 g/L n=3) of MDX. Doses of 5 g/L and 30 g/L MDX in the drinking water led to an extension of the lifespan in Ndufs4KO mice compared to vehicle (water)-treated mice. **B)** Survival curves of statistically significant MDX concentrations. * p<0.05, **p<0.01, Log-rank test vs vehicle.

### Maltodextrin administration does not improve clinical signs in Ndufs4KO mice

Ndufs4KO mice develop severe motor and breathing deficits as the pathology progresses (Quintana et al., 2010). Therefore, we set to assess whether the beneficial effects of MDX treatment (5 g/L or 30 g/L) in lifespan could also contribute to reduced Ndufs4KO symptomatology.

Ndufs4KO mice show reduced spontaneous locomotion and mobility (Chen et al., 2017). To assess the effect of MDX treatment on spontaneous locomotion in this mouse model animals treated with different doses of this compound were subjected to the open field (OF) test for 45 minutes, at early (P35), mid (P45) and late (P55) stage of the disease. During the first 5 minutes, corresponding to novelty-induced locomotion, Ndufs4KO mice moved significantly less than WT animals, regardless of the treatment, being already evident in MDX-treated mice at P35. However, no differences were observed between VEH-or MDX-treated Ndufs4KO mice (**Figure 3A**). In a similar fashion, analysis of the subsequent 40 minutes revealed significant differences between WT and Ndufs4KO mice, with no observable effect of MDX treatments in the performance of Ndufs4KO mice (**Figure 3B**).

**Figure 3.**
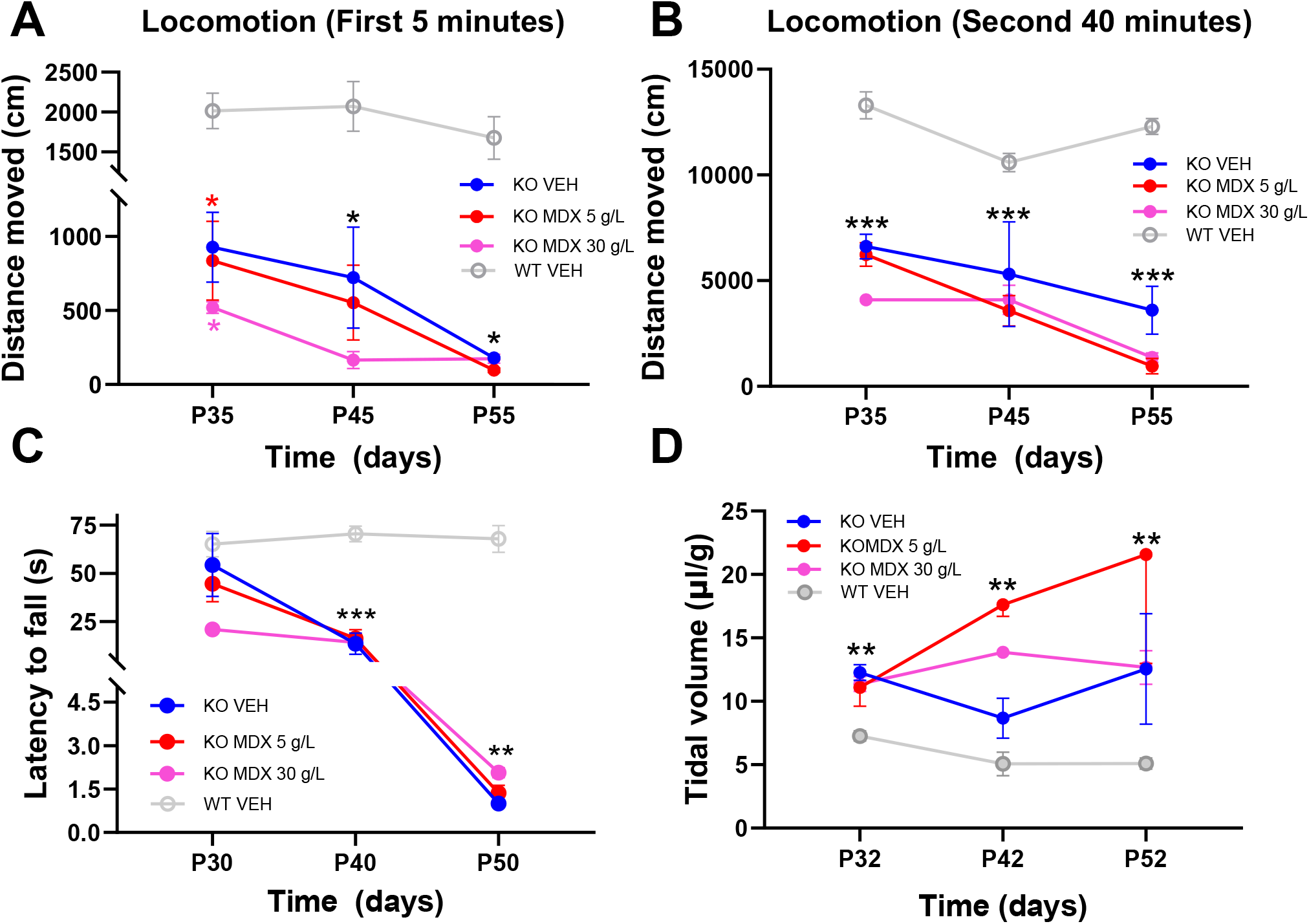
MDX administration does not correct motor or respiratory deficits in Ndufs4KO mice. **A-B)** Novelty-induced minute locomotion in WT or Ndufs4KO mice treated with either water (VEH) or 5 g/L or 30 g/L MDX in the drinking water during the first 5 minutes **(A)** or subsequent 40 minutes **(B)** in the open field test. **C)** Rotarod performance (latency to fall) in WT or Ndufs4KO mice treated with either water (VEH) or 5 g/L or 30 g/L MDX in the drinking water. **D)** Respiratory function (tidal volume) in WT or Ndufs4KO mice treated with either water (VEH) or 5 g/L or 30 g/L MDX in the drinking water * p<0.05, **p<0.01, ***p<0.001, mixed-model analysis. Black asterisks denote overall effect KO vs WT. Colored asterisks denote difference of the indicated group vs WT mice. n=3-6 per group.

Motor coordination was evaluated by the rotarod test at different stages of the disease. Animals lacking *Ndufs4* present impaired performance on the rotarod test after P40 (**Figure 3C**), indicating that motor coordination is affected in these animals, as described (Kruse et al., 2008; Sofou et al., 2014). No improvement was observed in animals treated with different doses of MDX when compared to vehicle-group, indicating that MDX has no effect on gross motor coordination in the Ndufs4KO mice in any stage of the disease (**Figure 3C**).

Respiratory alterations that have been linked to mortality both in Ndufs4KO mice (Quintana et al., 2012) and human patients (Lake et al., 2016). Specific respiratory parameters such as Tidal Volume (TV) are altered in the Ndufs4KO mice, accompanied with other breathing deficits such as hypo- and hyperventilation, gasping and apnea (Bolea et al., 2019). To assess the effect of MDX treatment on the respiratory function of Ndufs4KO mice, we analyzed TV by unrestrained whole-body plethysmography in awake mice at early, mid, and late (P52) stages of the disease **(Figure 3D)**, as described (Prada-Dacasa et al., 2020). In agreement with this, analysis of TV revealed significant differences in Ndufs4KO mice compared with WT mice. However, no significant differences were observed between VEH- and MDX-treated Ndufs4KO (**Figure 3D**). Thus, our results suggest that MDX treatment does not affect the development of clinical signs in Ndufs4KO mice.

### Maltodextrin administration reduces glial reactivity in Ndufs4KO mice

LS is characterized by symmetrical brain neuroinflammation followed by neurodegeneration in some brain nuclei, such us brainstem and basal ganglia in humans (Arii & Tanabe, 2000). In this regard, the Ndusf4KO mice the most affected area is the vestibular nucleus (VN) (Quintana et al., 2010).

To define the effect of MDX treatment on the overall neuroinflammatory phenotype in Ndufs4KO mice, immunofluorescence analysis for glial reactivity markers was performed in the VN of vehicle- and MDX-treated and MDX-treated (30 g/L) WT and Ndufs4KO mice (**Figure 4**). As expected, our results showed marked induction of astroglial marker glial fibrillary acidic protein (GFAP) and the microglial reactivity marker ionized calcium-binding adapter molecule 1 (Iba1) in the VN of VEH-treated Ndufs4KO mice compared to WT mice. However, this increase in glial reactivity was not present in MDX-treated Ndufs4KO mice (**Figure 4**), suggesting a decreased inflammatory tone in Ndufs4KO mice after MDX treatment.

**Figure 4.**
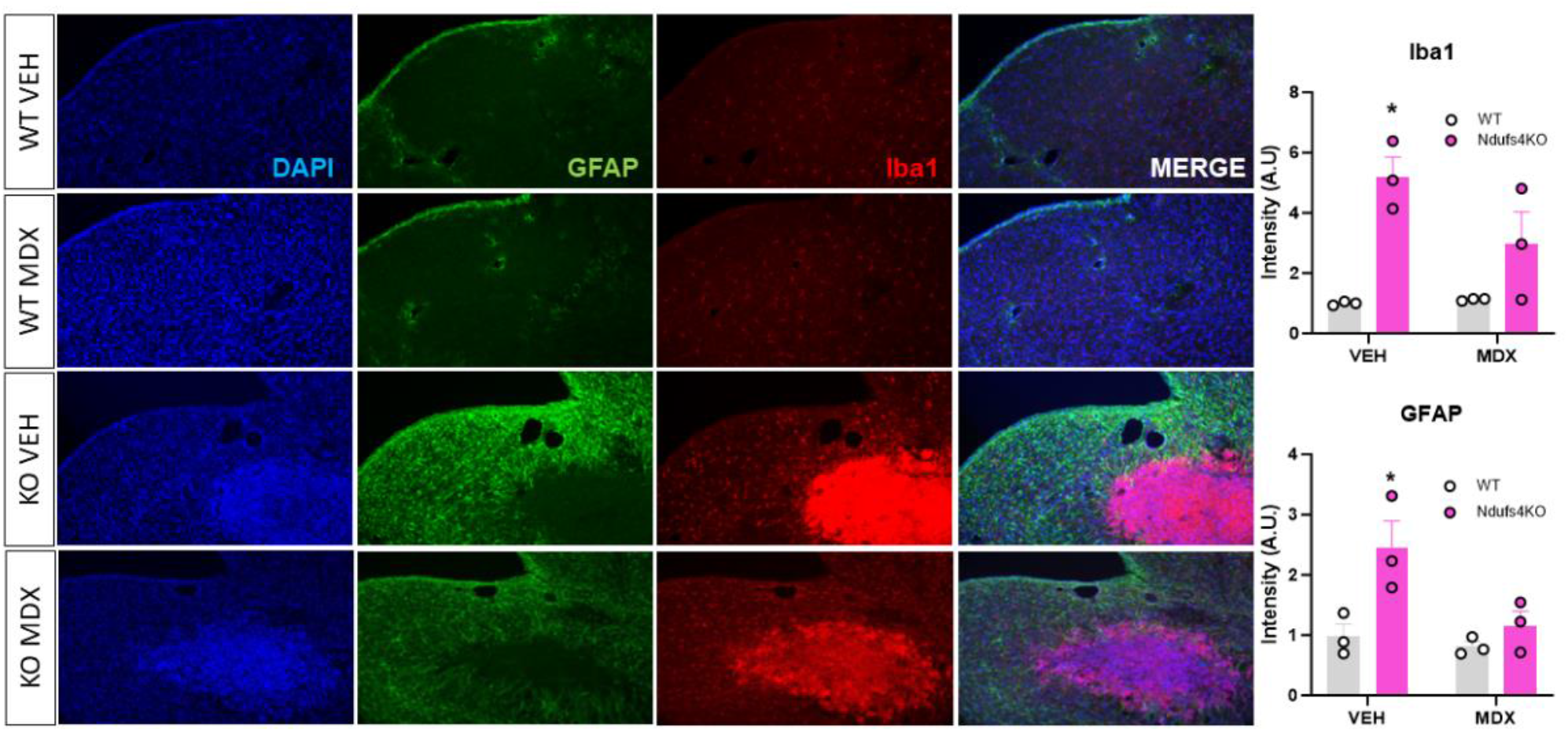
MDX administration reduces glial reactivity in the brainstem of Ndufs4KO mice. Immunofluorescence and quantification of the microglial marker Iba 1 (red), astroglial marker GFAP (green) in the vestibular nucleus region of VEH- and MDX-treated WT and Ndufs4KO mice. Nuclear counterstain (DAPI, blue) is also shown. * p<0.05 vs. WT VEH, two-way ANOVA Tukey’s multiple comparisons. n=3 per group.

### Altered intestinal microbiome profile in Ndufs4KO mice

We hypothesized that maltodextrin may be acting through regulation of intestinal microbiota in Ndufs4KO mice in a prebiotic manner. Thus, given the role that prebiotics play in modulating the host microbiota, high-throughput 16S rRNA gene sequencing of samples from the intestinal tract (caecum) was carried out in VEH- and MDX-treated (30 g/L) WT and Ndufs4KO mice. Metataxonomic analyses revealed a high variability between replicates within the same group, being thus very similar the resulting bacterial profiles among them. In terms of alpha-diversity (species richness), no significant differences were found (**Figure 5A**). In contrast, beta-diversity revealed a phenotype-dependent gut bacterial composition, but not a treatment dependent one (**Figure 5B**).

**Figure 5.**
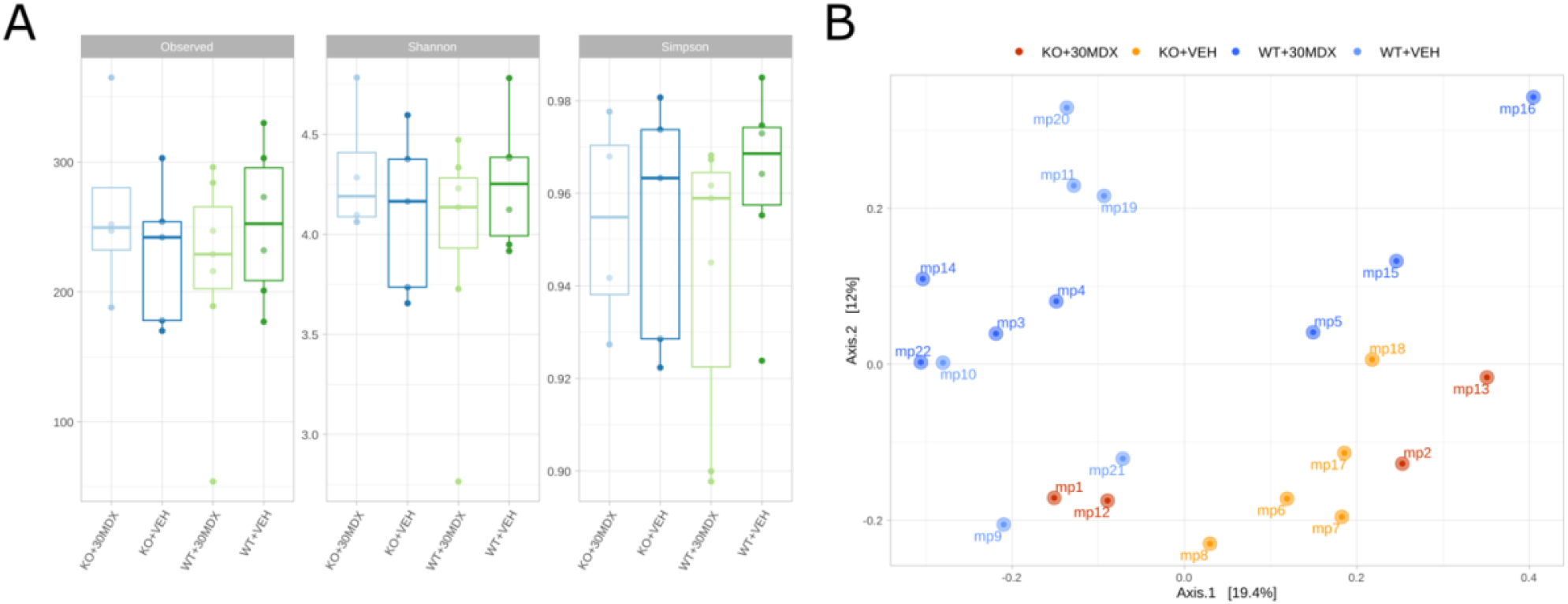
α- and β-diversity in caecal samples from VEH- and MDX-treated WT and Ndufs4KO mice. **(A)** α-diversity results at the ASV (Amplicon Sequence Variant) level. **(B)** β-diversity at the ASV level. n= 4-7per group.

No significant differences were observed at the phylum level among Ndufs4KO and WT mice (**Figure 6**). However, the comparison at the genus level (**Figure 7A-B**) revealed that the abundance of the genus *Akkermansia* was higher in Ndufs4KO mice (VEH) compared to WT (VEH) mice (**Figure 7B**). Furthermore, the abundance variability of *Akkermansia* was increased by MDX treatment, independent of the genotype (**Figure 7B**)

**Figure 6.**
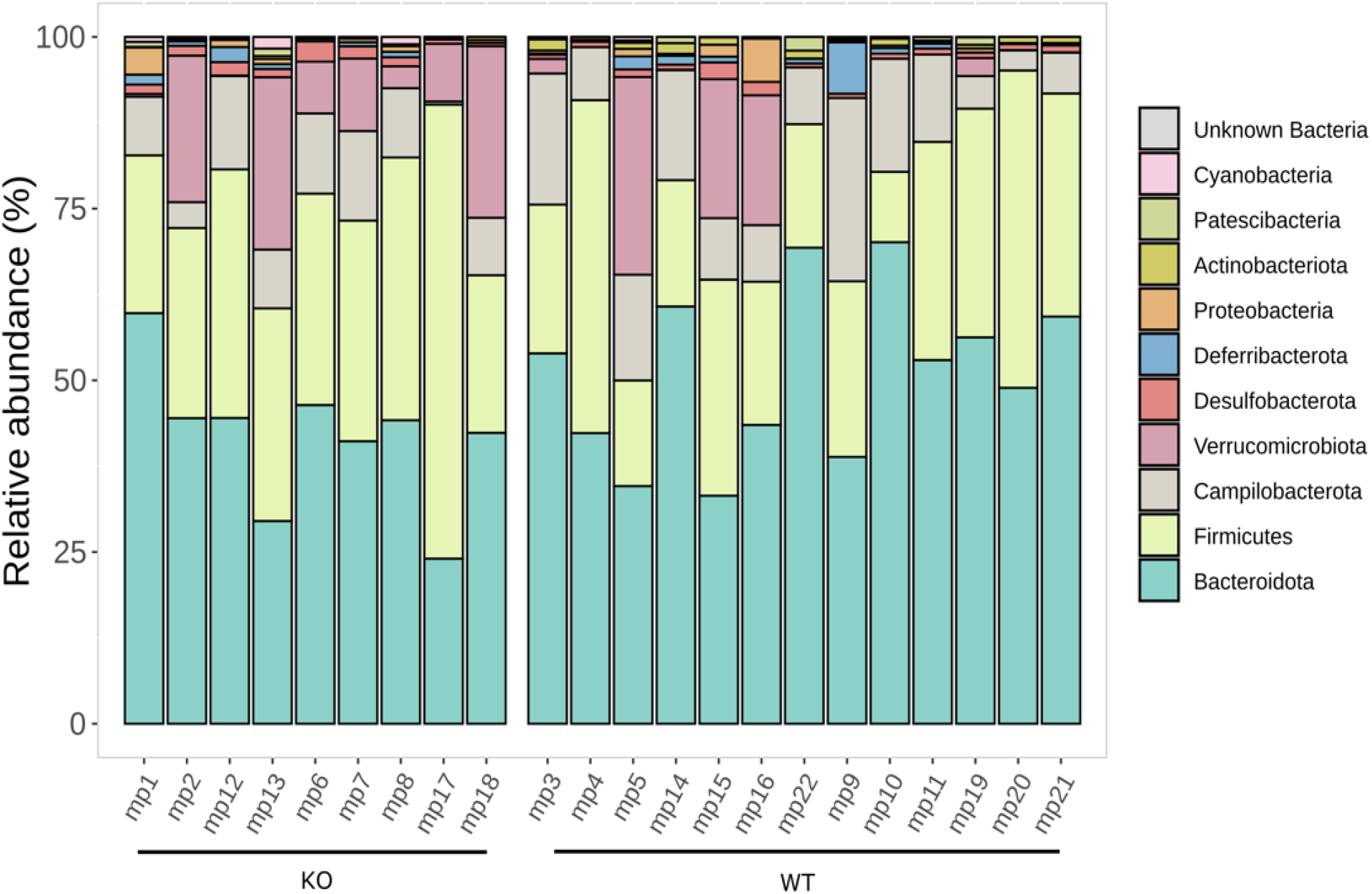
Bacterial profiles at the phylum level in caecal samples from VEH- and MDX-treated WT and Ndufs4KO mice. Similar profiles were obtained in all groups. n= 9-13 per group.

**Figure 7.**
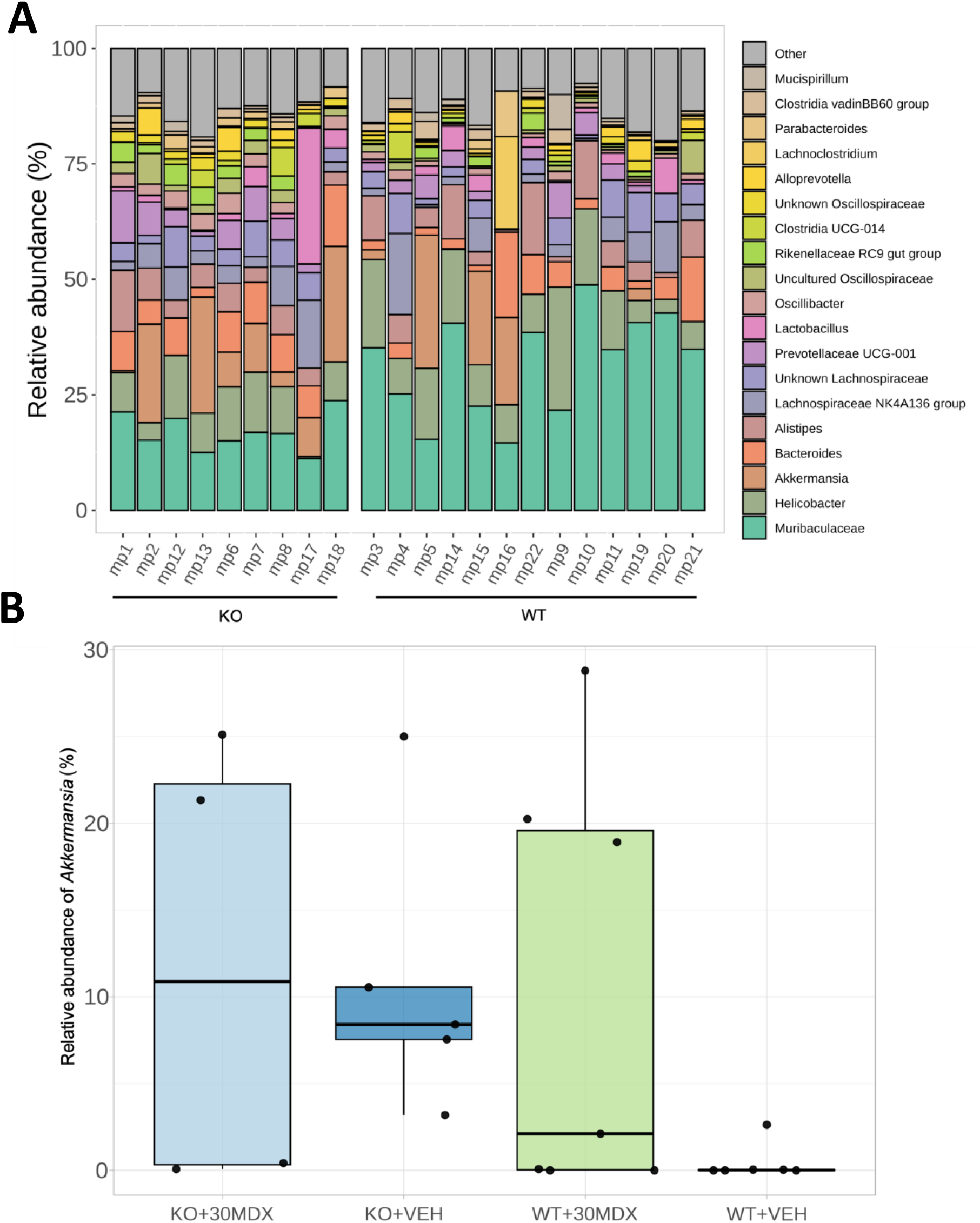
Bacterial profiles at the genus level in caecal samples from VEH- and MDX-treated WT and Ndufs4KO mice. **(A)** Global bacterial profiles at the genus level. **(B)** Relative abundances of *Akkermansia* (bottom) n= 4-7per group.

Independent of the treatment, group analysis between Ndufs4KO and WT mice was able to identify differences in several genera. In particular, NdufsKO mice presented increased abundance of *Biophila* and *Ruminococcus*. On the other hand, reduced abundance of *Muribaculaceae* was found in caeca from Ndufs4KO mice (**Figure 8**).

**Figure 8.**
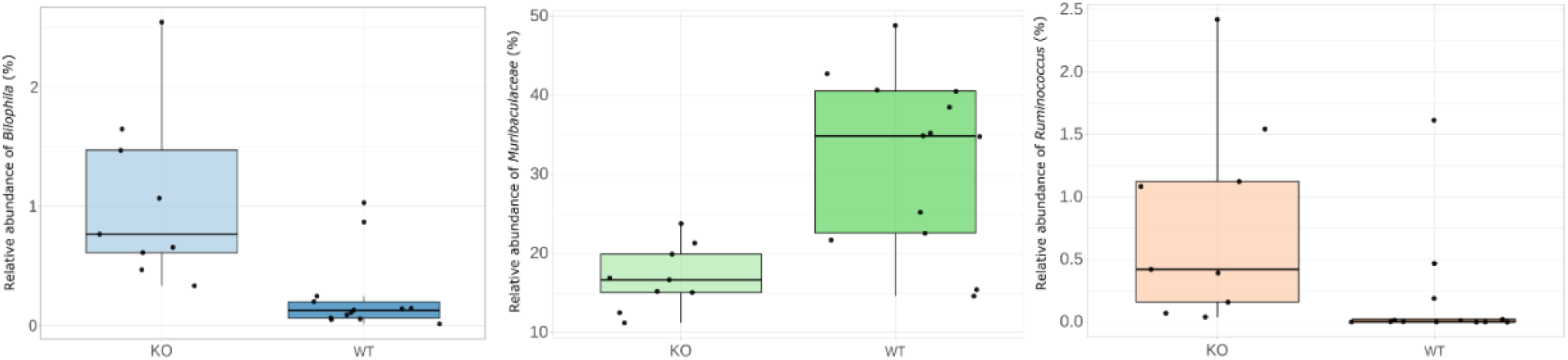
Relative abundance of *Bilophila* (left), *Muribaculaceae* (middle), and *Ruminococcus* (right) by phenotype. n= 4-7per group.

## Discussion

There is an urgent need to identify novel treatments for LS. Among the different treatment options, antioxidants have been shown to possess some value in animal models of Leigh Syndrome (de Haas et al., 2017; Liu et al., 2015). However, antioxidant approaches have had limited success in clinical settings. To that end, we hypothesized that high antioxidant loads may overcome this limitation, in line with previous studies (Liu et al., 2015). To that end, we set to use a high-antioxidant content microorganism, *Micrococcus luteus* (Molina-Menor et al., 2019), as a potential therapy for the Ndufs4KO (Quintana et al., 2010), a well-validated model of Leigh Syndrome. While we could not identify any beneficial effect attributed to *M. luteus* per se, here we show that, in a serendipitous discovery, the administration of maltodextrin (MDX), a cryopreservant used in the preparation of bacterial solutions, was sufficient to improve the phenotype of the Ndufs4KO mice.

MDX has many applications, mainly in food industry and as therapeutic approach in some pathologies related to nutrition and dehydration, where it has been demonstrated that is safe for human consumption (Hofman et al., 2016). However, the use of complex sugars for the treatment of LS or other MD had yet to be explored. Our results show a clear lifespan extension in Ndufs4KO mice treated with either 5 or 30 g/L of MDX, which was the highest dose test. Surprisingly, no effect on lifespan was observed with a dose of 15 g/L MDX. While this may be due to individual differences and/or statistical power, it could also underscore that the effect of MDX is not completely dependent on concentration and that may require the interaction with other factors. However, based on present results, we anticipate that the maximum dose-response effect on survival has not been reached. Therefore, further studies increasing the concentration of MDX will be necessary to define the dose leading to the maximal effect. In this regard, MDX at 30 g/L has a modest effect on the lifespan of Ndufs4KO mice when compared to another therapies for MD that have been tested in the same mouse model. Hypoxia (Ferrari et al., 2017), hyperoxia (Jain et al., 2019), rapamycin (Johnson et al., 2013),(Perry et al., 2021) and tetracyclines (Perry et al., 2021) treatment result in higher survival rates when compared to our results with MDX at 30 g/L in Ndufs4KO mice. However, we cannot exclude that higher doses of MDX may lead to improved effects on the lifespan of these animals.

The beneficial effects of MDX could be related to the capacity of MDX to keep Ndufs4KO mice hydrated for longer periods of time, and to the glucose from MDX molecule that provides the necessary energy for physiological processes to these animals. Here, it is important to point, that at late-stage of the disease, Ndufs4KO animals show severe locomotion alterations that hinder their capacity to drink and feed (Quintana et al., 2010).

Despite the encouraging results in lifespan extension in the MDX-treated Ndufs4KO mice, no clear effects were observed on the motor and respiratory alterations present in these animals. Analysis of locomotion in the open field test or motor coordination by rotarod test in the MDX-treated Ndufs4KO did not show any significant difference when compared to vehicle-treated group. However, the rotarod test evaluates gross coordination rather than fine motor coordination (Shiotsuki et al., 2010). For this reason, it remains to be determined whether MDX treatment affects other tests such as foot-print test and horizontal bar test, instead of rotarod which may be too demanding to appreciate differences in this severe phenotype.

Respiratory alterations are well described in Ndufs4KO mice and have been associated to increased lethality in both the Ndufs4KO model (Quintana et al., 2012) and LS patients (Arii & Tanabe, 2000). However, whole-body plethysmography analyzing TV, did not reveal significant differences between the MDX and vehicle-treated Ndufs4KO mice. In this regard, we and others have recently identified that spontaneous seizures are a driving cause of the lethal phenotype of the Ndufs4KO (Bolea et al., 2019; Bornstein et al., 2022) independent to motor and respiratory alterations. Thus, future studies to uncover whether MDX treatment ameliorates the epileptogenic phenotype are warranted.

Despite the lack of physiological and behavior deficits, MDX treatment is able to reduce the neuroinflammatory profile of Ndufs4KO mice in an affected area such as the VN. In this regard, MDX treatment at a concentration of 30 g/L reduces astrocyte and microglia-containing lesions in the VN of Ndufs4KO mice when compared to vehicle-treated Ndufs4KO mice. In this regard, increased glial reactivity has been associated with poor outcome and death in Ndufs4KO mice (Quintana et al., 2012). In line with this, interventions leading to reduced inflammation (Johnson et al., 2013; Liu et al., 2015; Perry et al., 2021) or targeting immune cells (Aguilar et al., 2022; Stokes et al., 2022) have led to increased lifespan in the Ndufs4KO mice. Thus, it is likely that the reduction of glial activity is playing a role in the beneficial effects of MDX treatment.

However, it is still unclear how MDX may exert its beneficial effects. It has been described that MDX possesses antioxidant capabilities (Hofman et al., 2016). However, it is likely that this effect may be locally observed in the gut, rather than the brain. Hence, given the known and growing relationship between microbiota, brain and inflammation (Rutsch et al., 2020), we hypothesized that MDX may be affecting microbiota in the intestines of Ndufs4KO mice in a prebiotic manner. In this regard, even though interindividual variability has limited the ability to identify differences at the phyla level, our metataxonomic analyses have identified several genera altered in the intestines of Ndufs4KO mice compared to WT. Among them, we have detected an increase in the relative abundance of *Bilophila* and *Ruminococcus*, coincident with reports in neurodegenerative diseases such as Parkinson Disease (Li et al., 2023). Along the same lines, *Muribaculaceae* has been suggested to negatively correlate with inflammation (Shang et al., 2021). Thus, the alterations observed in intestinal microbiota in Ndufs4KO mice seem to point to an enhanced inflammatory status in these mice, which may contribute to pathology progression, in line with recent reports (Bitto et al., 2023). As mentioned above, no robust effects were found in bacterial profiling after MDX administration. However, *Akkermansia* was significantly increased in Ndufs4KO VEH mice compared to WT VEH controls. The role of *Akkermansia* in physiology and pathology is gaining traction, having been described both positive and negative roles in inflammation and pathological progression in different paradigms (Wang et al., 2022; Zhang et al., 2019). MDX treatment, both in Ndufs4KO and WT mice, was associated with an increase in the variability of *Akkermansia* abundance, which could underlie some of the beneficial effects of MDX. In this regard, further in-depth studies to identify possible correlations between microbiome profiles, *Akkermansia* abundance, MDX treatment, and lifespan extension can help identify the underlying factors driving the beneficial effects of MDX administration in Ndusf4KO.

In conclusion, our results pave the way for the development of MDX as a potential novel treatment for Leigh Syndrome, and identify alterations in the intestinal microbiota in Ndus4KO that may be involved in the progression of the disease.

## Author contributions

ADM, EMM, MBR and AU carried out the experiments, and generated and analyzed data. JP, MP and AQ planned, designed, and supervised the experiments and obtained resources. AMD, EMM and AQ wrote the manuscript.

## Acknowledgments

This work was supported by a MINECO Ramon y Cajal fellowships (RyC-2012-11873; A.Q.), and pre-doctoral fellowships (Formación de Doctores PRE2021-096929 to ADM, AGAUR Joan Oró 2023 FI-100663 to MBR, 2018FI_B 00452 to AU, Formación del Profesorado Universitario grant with reference FPU17/04184 to EMM). A.Q. received funds from the European Research Council (Starting grant NEUROMITO, ERC-2014-StG-638106), MINECO Proyectos I+D de Excelencia (SAF2014-57981P; SAF2017-88108-R), MICINN Proyectos I+D+i (PID2020-114977RB-I00), AGAUR (2017SGR-323), Fundació TV3-La Marató (202030), and “la Caixa” Foundation (ID 100010434), under the agreement LCF/PR/HR20/52400018. We acknowledge Adriel Latorre-Pérez for his assistance and help with metataxonomy and Darwin Bioprospecting Excellence S.L. for generously contributing to this work for free.

## References

Aguilar, K., Comes, G., Canal, C., Quintana, A., Sanz, E., & Hidalgo, J. (2022). Microglial response promotes neurodegeneration in the Ndufs4 KO mouse model of Leigh syndrome. Glia, 70(11), 2032–2044. https://doi.org/10.1002/glia.24234

Angelova, P. R., & Abramov, A. Y. (2018). Role of mitochondrial ROS in the brain: from physiology to neurodegeneration. FEBS Lett, 592(5), 692–702. https://doi.org/10.1002/1873-3468.12964

Apel, K., & Hirt, H. (2004). Reactive oxygen species: metabolism, oxidative stress, and signal transduction. Annu Rev Plant Biol, 55, 373–399. https://doi.org/10.1146/annurev.arplant.55.031903.141701

Arii, J., & Tanabe, Y. (2000). Leigh syndrome: serial MR imaging and clinical follow-up. AJNR Am J Neuroradiol, 21(8), 1502–1509. https://www.ncbi.nlm.nih.gov/pubmed/11003287

Asker, D., Beppu, T., & Ueda, K. (2007). Unique diversity of carotenoid-producing bacteria isolated from Misasa, a radioactive site in Japan. Appl Microbiol Biotechnol, 77(2), 383–392. https://doi.org/10.1007/s00253-007-1157-8

Bitto, A., Grillo, A. S., Ito, T. K., Stanaway, I. B., Nguyen, B. M. G., Ying, K., Tung, H., Smith, K., Tran, N., Velikanje, G., Urfer, S. R., Snyder, J. M., Barton, J., Sharma, A., Kayser, E. B., Wang, L., Smith, D. L., Jr., Thompson, J. W., DuBois, L., Kaeberlein, M. (2023). Acarbose suppresses symptoms of mitochondrial disease in a mouse model of Leigh syndrome. Nat Metab, 5(6), 955–967. https://doi.org/10.1038/s42255-023-00815-w

Bolea, I., Gella, A., Sanz, E., Prada-Dacasa, P., Menardy, F., Bard, A. M., Machuca-Marquez, P., Eraso-Pichot, A., Modol-Caballero, G., Navarro, X., Kalume, F., & Quintana, A. (2019). Defined neuronal populations drive fatal phenotype in a mouse model of Leigh syndrome. Elife, 8. https://doi.org/10.7554/eLife.47163

Bornstein, R., James, K., Stokes, J., Park, K. Y., Kayser, E. B., Snell, J., Bard, A., Chen, Y., Kalume, F., & Johnson, S. C. (2022). Differential effects of mTOR inhibition and dietary ketosis in a mouse model of subacute necrotizing encephalomyelopathy. Neurobiol Dis, 163, 105594. https://doi.org/10.1016/j.nbd.2021.105594

Chen, B., Hui, J., Montgomery, K. S., Gella, A., Bolea, I., Sanz, E., Palmiter, R. D., & Quintana, A. (2017). Loss of Mitochondrial Ndufs4 in Striatal Medium Spiny Neurons Mediates Progressive Motor Impairment in a Mouse Model of Leigh Syndrome. Front Mol Neurosci, 10, 265. https://doi.org/10.3389/fnmol.2017.00265

Cobb, C. A., & Cole, M. P. (2015). Oxidative and nitrative stress in neurodegeneration. Neurobiol Dis, 84, 4–21. https://doi.org/10.1016/j.nbd.2015.04.020

Cunningham, M., Azcarate-Peril, M. A., Barnard, A., Benoit, V., Grimaldi, R., Guyonnet, D., Holscher, H. D., Hunter, K., Manurung, S., Obis, D., Petrova, M. I., Steinert, R. E., Swanson, K. S., van Sinderen, D., Vulevic, J., & Gibson, G. R. (2021). Shaping the Future of Probiotics and Prebiotics. Trends Microbiol, 29(8), 667–685. https://doi.org/10.1016/j.tim.2021.01.003

de Haas, R., Das, D., Garanto, A., Renkema, H. G., Greupink, R., van den Broek, P., Pertijs, J., Collin, R. W. J., Willems, P., Beyrath, J., Heerschap, A., Russel, F. G., & Smeitink, J. A. (2017). Therapeutic effects of the mitochondrial ROS-redox modulator KH176 in a mammalian model of Leigh Disease. Sci Rep, 7(1), 11733. https://doi.org/10.1038/s41598-017-09417-5

Dumitrescu, L., Popescu-Olaru, I., Cozma, L., Tulba, D., Hinescu, M. E., Ceafalan, L. C., Gherghiceanu, M., & Popescu, B. O. (2018). Oxidative Stress and the Microbiota-Gut-Brain Axis. Oxid Med Cell Longev, 2018, 2406594. https://doi.org/10.1155/2018/2406594

Enns, G. M. (2014). Treatment of mitochondrial disorders: antioxidants and beyond. J Child Neurol, 29(9), 1235–1240. https://doi.org/10.1177/0883073814538509

Fan, Y., & Pedersen, O. (2021). Gut microbiota in human metabolic health and disease. Nat Rev Microbiol, 19(1), 55–71. https://doi.org/10.1038/s41579-020-0433-9

Ferrari, M., Jain, I. H., Goldberger, O., Rezoagli, E., Thoonen, R., Cheng, K. H., Sosnovik, D. E., Scherrer-Crosbie, M., Mootha, V. K., & Zapol, W. M. (2017). Hypoxia treatment reverses neurodegenerative disease in a mouse model of Leigh syndrome. Proc Natl Acad Sci U S A, 114(21), E4241–E4250. https://doi.org/10.1073/pnas.1621511114

Friedman, J. R., & Nunnari, J. (2014). Mitochondrial form and function. Nature, 505(7483), 335–343. https://doi.org/10.1038/nature12985

Gerards, M., Sallevelt, S. C., & Smeets, H. J. (2016). Leigh syndrome: Resolving the clinical and genetic heterogeneity paves the way for treatment options. Mol Genet Metab, 117(3), 300–312. https://doi.org/10.1016/j.ymgme.2015.12.004

Grimm, A., & Eckert, A. (2017). Brain aging and neurodegeneration: from a mitochondrial point of view. J Neurochem, 143(4), 418–431. https://doi.org/10.1111/jnc.14037

Hofman, D. L., van Buul, V. J., & Brouns, F. J. (2016). Nutrition, Health, and Regulatory Aspects of Digestible Maltodextrins. Crit Rev Food Sci Nutr, 56(12), 2091–2100. https://doi.org/10.1080/10408398.2014.940415

Jain, I. H., Zazzeron, L., Goldberger, O., Marutani, E., Wojtkiewicz, G. R., Ast, T., Wang, H., Schleifer, G., Stepanova, A., Brepoels, K., Schoonjans, L., Carmeliet, P., Galkin, A., Ichinose, F., Zapol, W. M., & Mootha, V. K. (2019). Leigh Syndrome Mouse Model Can Be Rescued by Interventions that Normalize Brain Hyperoxia, but Not HIF Activation. Cell Metab, 30(4), 824–832 e823. https://doi.org/10.1016/j.cmet.2019.07.006

Johnson, S. C., Yanos, M. E., Kayser, E. B., Quintana, A., Sangesland, M., Castanza, A., Uhde, L., Hui, J., Wall, V. Z., Gagnidze, A., Oh, K., Wasko, B. M., Ramos, F. J., Palmiter, R. D., Rabinovitch, P. S., Morgan, P. G., Sedensky, M. M., & Kaeberlein, M. (2013). mTOR inhibition alleviates mitochondrial disease in a mouse model of Leigh syndrome. Science, 342(6165), 1524–1528. https://doi.org/10.1126/science.1244360

Kruse, S. E., Watt, W. C., Marcinek, D. J., Kapur, R. P., Schenkman, K. A., & Palmiter, R. D. (2008). Mice with mitochondrial complex I deficiency develop a fatal encephalomyopathy. Cell Metab, 7(4), 312–320. https://doi.org/10.1016/j.cmet.2008.02.004

Lake, N. J., Compton, A. G., Rahman, S., & Thorburn, D. R. (2016). Leigh syndrome: One disorder, more than 75 monogenic causes. Ann Neurol, 79(2), 190–203. https://doi.org/10.1002/ana.24551

Lavie, L. (2015). Oxidative stress in obstructive sleep apnea and intermittent hypoxia--revisited--the bad ugly and good: implications to the heart and brain. Sleep Med Rev, 20, 27–45. https://doi.org/10.1016/j.smrv.2014.07.003

Li, Z., Liang, H., Hu, Y., Lu, L., Zheng, C., Fan, Y., Wu, B., Zou, T., Luo, X., Zhang, X., Zeng, Y., Liu, Z., Zhou, Z., Yue, Z., Ren, Y., Li, Z., Su, Q., & Xu, P. (2023). Gut bacterial profiles in Parkinson’s disease: A systematic review. CNS Neurosci Ther, 29(1), 140–157. https://doi.org/10.1111/cns.13990

Liu, L., Zhang, K., Sandoval, H., Yamamoto, S., Jaiswal, M., Sanz, E., Li, Z., Hui, J., Graham, B. H., Quintana, A., & Bellen, H. J. (2015). Glial lipid droplets and ROS induced by mitochondrial defects promote neurodegeneration. Cell, 160(1-2), 177–190. https://doi.org/10.1016/j.cell.2014.12.019

Lynch, S. V., & Pedersen, O. (2016). The Human Intestinal Microbiome in Health and Disease. N Engl J Med, 375(24), 2369–2379. https://doi.org/10.1056/NEJMra1600266

Martinelli, D., Catteruccia, M., Piemonte, F., Pastore, A., Tozzi, G., Dionisi-Vici, C., Pontrelli, G., Corsetti, T., Livadiotti, S., Kheifets, V., Hinman, A., Shrader, W. D., Thoolen, M., Klein, M. B., Bertini, E., & Miller, G. (2012). EPI-743 reverses the progression of the pediatric mitochondrial disease--genetically defined Leigh Syndrome. Mol Genet Metab, 107(3), 383–388. https://doi.org/10.1016/j.ymgme.2012.09.007

Milani, A., Basirnejad, M., Shahbazi, S., & Bolhassani, A. (2017). Carotenoids: biochemistry, pharmacology and treatment. Br J Pharmacol, 174(11), 1290–1324. https://doi.org/10.1111/bph.13625

Molina-Menor, E., Gimeno-Valero, H., Pascual, J., Pereto, J., & Porcar, M. (2020). High Culturable Bacterial Diversity From a European Desert: The Tabernas Desert. Front Microbiol, 11, 583120. https://doi.org/10.3389/fmicb.2020.583120

Molina-Menor, E., Tanner, K., Vidal-Verdu, A., Pereto, J., & Porcar, M. (2019). Microbial communities of the Mediterranean rocky shore: ecology and biotechnological potential of the sea-land transition. Microb Biotechnol, 12(6), 1359–1370. https://doi.org/10.1111/1751-7915.13475

Ng, Y. S., & Turnbull, D. M. (2016). Mitochondrial disease: genetics and management. J Neurol, 263(1), 179–191. https://doi.org/10.1007/s00415-015-7884-3

Parkinson, M. H., Schulz, J. B., & Giunti, P. (2013). Co-enzyme Q10 and idebenone use in Friedreich’s ataxia. J Neurochem, 126 Suppl 1, 125–141. https://doi.org/10.1111/jnc.12322

Perry, E. A., Bennett, C. F., Luo, C., Balsa, E., Jedrychowski, M., O’Malley, K. E., Latorre-Muro, P., Ladley, R. P., Reda, K., Wright, P. M., Gygi, S. P., Myers, A. G., & Puigserver, P. (2021). Tetracyclines promote survival and fitness in mitochondrial disease models. Nat Metab, 3(1), 33–42. https://doi.org/10.1038/s42255-020-00334-y

Prada-Dacasa, P., Urpi, A., Sanchez-Benito, L., Bianchi, P., & Quintana, A. (2020). Measuring Breathing Patterns in Mice Using Whole-body Plethysmography. Bio Protoc, 10(17), e3741. https://doi.org/10.21769/BioProtoc.3741

Quintana, A., Kruse, S. E., Kapur, R. P., Sanz, E., & Palmiter, R. D. (2010). Complex I deficiency due to loss of Ndufs4 in the brain results in progressive encephalopathy resembling Leigh syndrome. Proc Natl Acad Sci U S A, 107(24), 10996–11001. https://doi.org/10.1073/pnas.1006214107

Quintana, A., Zanella, S., Koch, H., Kruse, S. E., Lee, D., Ramirez, J. M., & Palmiter, R. D. (2012). Fatal breathing dysfunction in a mouse model of Leigh syndrome. J Clin Invest, 122(7), 2359–2368. https://doi.org/10.1172/JCI62923

Rahman, S. (2012). Mitochondrial disease and epilepsy. Dev Med Child Neurol, 54(5), 397–406. https://doi.org/10.1111/j.1469-8749.2011.04214.x

Rutsch, A., Kantsjo, J. B., & Ronchi, F. (2020). The Gut-Brain Axis: How Microbiota and Host Inflammasome Influence Brain Physiology and Pathology. Front Immunol, 11, 604179. https://doi.org/10.3389/fimmu.2020.604179

Shang, L., Liu, H., Yu, H., Chen, M., Yang, T., Zeng, X., & Qiao, S. (2021). Core Altered Microorganisms in Colitis Mouse Model: A Comprehensive Time-Point and Fecal Microbiota Transplantation Analysis. Antibiotics (Basel), 10(6). https://doi.org/10.3390/antibiotics10060643

Shiotsuki, H., Yoshimi, K., Shimo, Y., Funayama, M., Takamatsu, Y., Ikeda, K., Takahashi, R., Kitazawa, S., & Hattori, N. (2010). A rotarod test for evaluation of motor skill learning. J Neurosci Methods, 189(2), 180–185. https://doi.org/10.1016/j.jneumeth.2010.03.026

Sofou, K., De Coo, I. F., Isohanni, P., Ostergaard, E., Naess, K., De Meirleir, L., Tzoulis, C., Uusimaa, J., De Angst, I. B., Lonnqvist, T., Pihko, H., Mankinen, K., Bindoff, L. A., Tulinius, M., & Darin, N. (2014). A multicenter study on Leigh syndrome: disease course and predictors of survival. Orphanet J Rare Dis, 9, 52. https://doi.org/10.1186/1750-1172-9-52

Soheili, M., Alinaghipour, A., & Salami, M. (2022). Good bacteria, oxidative stress and neurological disorders: Possible therapeutical considerations. Life Sci, 301, 120605. https://doi.org/10.1016/j.lfs.2022.120605

Stahl, W., & Sies, H. (2005). Bioactivity and protective effects of natural carotenoids. Biochim Biophys Acta, 1740(2), 101–107. https://doi.org/10.1016/j.bbadis.2004.12.006

Stokes, J. C., Bornstein, R. L., James, K., Park, K. Y., Spencer, K. A., Vo, K., Snell, J. C., Johnson, B. M., Morgan, P. G., Sedensky, M. M., Baertsch, N. A., & Johnson, S. C. (2022). Leukocytes mediate disease pathogenesis in the Ndufs4(KO) mouse model of Leigh syndrome. JCI Insight, 7(5). https://doi.org/10.1172/jci.insight.156522

Tanner, K., Martorell, P., Genoves, S., Ramon, D., Zacarias, L., Rodrigo, M. J., Pereto, J., & Porcar, M. (2019). Bioprospecting the Solar Panel Microbiome: High-Throughput Screening for Antioxidant Bacteria in a Caenorhabditis elegans Model. Front Microbiol, 10, 986. https://doi.org/10.3389/fmicb.2019.00986

Tapia, C., Lopez, B., Astuya, A., Becerra, J., Gugliandolo, C., Parra, B., & Martinez, M. (2021). Antiproliferative activity of carotenoid pigments produced by extremophile bacteria. Nat Prod Res, 35(22), 4638–4642. https://doi.org/10.1080/14786419.2019.1698574

Tian, B., & Hua, Y. (2010). Carotenoid biosynthesis in extremophilic Deinococcus-Thermus bacteria. Trends Microbiol, 18(11), 512–520. https://doi.org/10.1016/j.tim.2010.07.007

Vafai, S. B., & Mootha, V. K. (2012). Mitochondrial disorders as windows into an ancient organelle. Nature, 491(7424), 374–383. https://doi.org/10.1038/nature11707

Wang, K., Wu, W., Wang, Q., Yang, L., Bian, X., Jiang, X., Lv, L., Yan, R., Xia, J., Han, S., & Li, L. (2022). The negative effect of Akkermansia muciniphila-mediated post-antibiotic reconstitution of the gut microbiota on the development of colitis-associated colorectal cancer in mice. Front Microbiol, 13, 932047. https://doi.org/10.3389/fmicb.2022.932047

Zhang, T., Li, Q., Cheng, L., Buch, H., & Zhang, F. (2019). Akkermansia muciniphila is a promising probiotic. Microb Biotechnol, 12(6), 1109–1125. https://doi.org/10.1111/1751-7915.13410

